# Negros Bleeding-heart *Gallicolumba keayi* prefers dense understorey vegetation and dense canopy cover, and species distribution modelling shows little remaining suitable habitat

**DOI:** 10.1101/872044

**Authors:** Holly Mynott, Mark Abrahams, Daphne Kerhoas

## Abstract

The Philippines is a global biodiversity hotspot, with a large number of Threatened bird species, one of which is the Critically Endangered Negros Bleeding-heart *Gallicolumba keayi*. The aim of this study was to investigate the habitat preference of the Negros Bleeding-heart and undertake species distribution modelling to locate areas of conservation importance based on identified suitable habitat. A survey of 94 point counts was undertaken and eight camera traps were deployed from May to August 2018 in the Northwest Panay Peninsula Natural Park, Panay, Philippines. Habitat variables (canopy cover, understorey cover, ground cover, altitude, presence of rattan and pandan, tree diameter at breast height and branching architecture) were measured in 93 5 m-radius quadrats. To identify areas of potentially suitable habitat for the Negros Bleeding-heart, species distribution modelling was undertaken in MaxEnt using tree cover and altitude data on Panay and Negros. Using a Generalised Linear Model, Negros Bleeding-heart presence was found to be significantly positively associated with high understorey cover and dense canopy cover. Species distribution modelling showed that the Northwest Panay Peninsula Natural Park is currently the most effectively located protected area for Negros Bleeding-heart conservation, while protected areas in Negros require further protection. It is imperative that protection is continued in the Northwest Panay Peninsula Natural Park, and more survey effort is needed to identify other critical Negros Bleeding-heart populations, around which deforestation and hunting ban enforcement is strongly recommended.

## Introduction

The Philippines is one of the top five global biodiversity hotspots, with 46% species endemism (Lee *et al.* 2012). However, forest cover has fallen from 90% to 27% from 1521 to the present, and only 11% of what remains is primary forest (Food and Agriculture Organisation of the United Nations 2015). This has been a contributing factor to bird species declines, and the country now has the 5^th^ highest number of Critically Endangered bird species in the world (BirdLife International 2008a). It is therefore a priority area for global conservation (Paz *et al.* 2013).

The Negros Bleeding-heart is a Critically Endangered colombid, found only on the islands of Panay and Negros in the Philippines. The most recent population estimate is from 2001, when 70 – 370 individuals were thought to remain in the wild (BirdLife International 2017). It is threatened by habitat loss through forest conversion to agriculture or clearance for timber extraction and charcoal production; from the 1500s to 1988, forest cover on Panay and Negros was reduced by 92% and 96% respectively, and very little primary forest remains (BirdLife International 2017). It is also threatened by poaching, as it is trapped, and hunted for food and to keep as a pet (Cariño 2007; BirdLife International 2017). Very few conservation measures are in place apart from patrols by forest rangers to dismantle traps and stop illegal logging, e.g. in the Northwest Panay Peninsula Natural Park and northern Central Panay Mountain Range (Arkive 2007). Due to its small population and ongoing threats, the Negros Bleeding-heart urgently requires conservation action (Bristol Zoo Gardens 2017; Gaworecki 2018).

In terms of its biology and resource requirements, much remains unknown about the Negros Bleeding-heart, as is the case for most species in the *Gallicolumba* family (Walker 2007). It is a ground-dwelling bird that spends most of its time on the forest floor and only uses understorey plants to roost, take cover, or breed (Slade *et al.* 2005). It is cryptic and difficult to observe (Slade *et al.* 2005; Cariño 2007). Its diet consists of seeds, berries and invertebrates (Slade *et al.* 2005; Cariño 2007), including fruit from *Ficus* and *Pinanga* species (Cariño 2007). It is thought to prefer lowland forest 300-1000 m (BirdLife International 2017) but it has also been reported >1000 m on Negros, albeit where all forest <800 m has been cleared (Curio 2001). It is thought to prefer primary forest, but its ability to utlise secondary forest remains unclear; although reportedly detected in secondary forest on Panay, it has not been recorded in secondary forest on Negros (BirdLife International 2017). Understanding more about its resource requirements and habitat preference is important for its conservation (Bibby *et al.* 2000; Begehold *et al.* 2015). It can be used in species distribution modelling, in which species presence and environmental data is used to map where a species is most probably distributed (Morales *et al.* 2017), and this is used to identify priority areas for conservation action, based on an evaluation of what already exists and what is needed (De Carvalho *et al.* 2017).

The most recent study of the Negros Bleeding-heart was commissioned by Bristol Zoological Society and undertaken by the Center for Conservation Innovation (CCI), investigating its population size and habitat preference in south Negros (CCI report, personal communication). Based on nine observations in March 2016, occupancy modelling found that the dove prefers sites with dense ground vegetation coverage and high canopy cover (CCI report, personal communication). Furthermore, bird-habitat association modelling suggested an association with ground cover, midstorey cover, fruiting and flowering trees, and large trees of 20-60 cm diameter at breast height (CCI report, personal communication).

The aim of the current study was to investigate the habitat preference of the Negros Bleeding-heart in the Northwest Panay Peninsula Natural Park and to model species distribution across its range to identify the best potential areas for conservation action. Objectives therefore were to (1) survey for the Negros Bleeding-heart using point counts, line transects, playback surveys and camera traps and collect habitat variable measurements, and assess the influence of measured habitat variables on this species’ presence using Generalised Linear Modelling (GLM); and (2) identify areas of potentially suitable habitat across Panay and Negros using species distribution modelling, and compare Negros Bleeding-heart predicted distribution against current protected areas and Important Bird Areas to identify focus areas for future conservation action.

## Methods

### Data Collection

#### Study site

The study was undertaken in the forest surrounding Sibaliw field station (latitude 121.9675, longitude 11.8195) in the Northwest Panay Peninsula Natural Park, Panay, Philippines (figure 1). At 5,000 ha, at least half of which is old growth, this encompasses the largest remaining low elevation forest across Negros and Panay (BirdLife International 2018a). It is a protected area, where hunting, trapping or disturbing wild plants or animals, cutting timber or gathering forest products and shifting cultivation are prohibited (Zabal 2014). Various Endangered species have been observed there, such as the Visayan warty pig *Sus cebifrons* (CR) (Meijaard *et al.* 2017), Visayan Hornbill *Penelopides panini* (EN), Negros Bleeding-heart *Gallicolumba keayi* (CR) (BirdLife International 2018a), White-throated Jungle Flycatcher *Vauriella albigularis* (EN) and Yellow-faced Flameback *Chrysocolaptes xanthocephalus* (EN) (Kennedy *et al.* 2000). Deforestation, mining, hunting and poaching have also been observed inside the protected area (Foundation for the Philippine Environment 2018).

**Figure 1.**
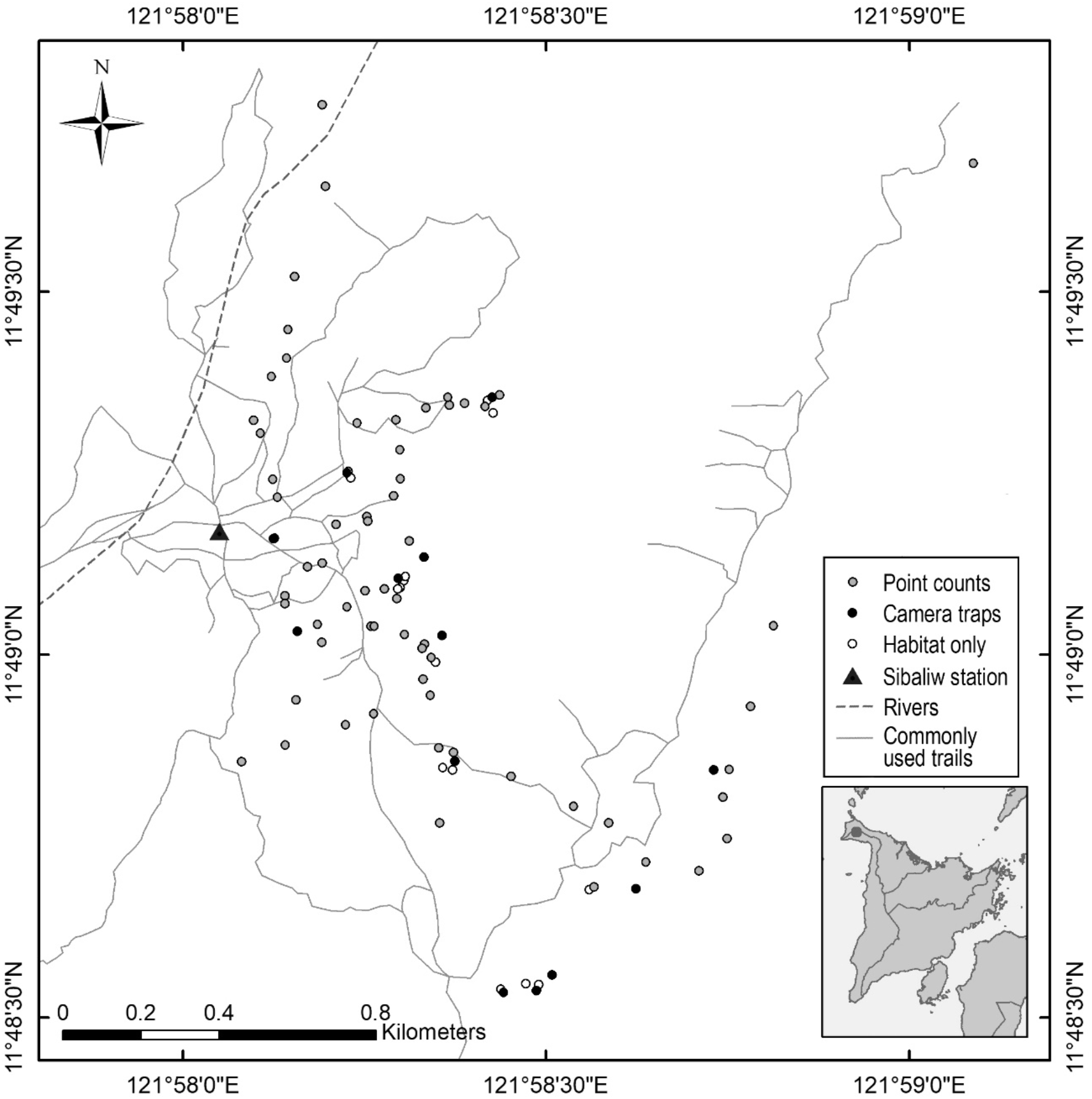
The location of the study site, within the Northwest Panay Peninsula Natural Park, Panay, Philippines. Commonly used trails and points at which data was collected (circles) are mapped. Rivers (OCHA Philippines 2013) are also shown. Map inset shows the location of Sibaliw research station on the island of Panay, Philippines.

#### Surveying for birds

Two expeditions were undertaken in the same location within the Northwest Panay Peninsula Natural Park to survey for the Negros Bleeding-heart. Due to the extreme rarity of the species, all contacts with it, obtained using different methods, were used in this analysis to obtain information about distribution and habitat association. The first expedition was from 9-22 May 2018 (total 13 days observation) by Mark Abrahams, Daphne Kerhoas and Jenny Poole from Bristol Zoological Society with guides Potpot Fernandez and Jun Tacud. 20 line transects of 500 m length were undertaken, with 30 point counts at the start and at the end of each transect. Point counts (where an observer records all birds seen and heard from one location; Lloyd *et al.* 2000) lasted for 15 minutes, including a 5 minute settling period. Sightings of Negros Bleeding-heart were also recorded *ad libitum*, and three playback surveys were undertaken. The second survey was from 12 July – 6 August 2018 (total 25 days observation) by Holly Mynott and the same two guides, in which 64 point counts that were 10 minutes long were undertaken as recommended for cryptic (Bibby *et al.* 2000) and terrestrial (Lee and Marsden 2008) bird species. This totals 94 point counts, of which 80 were laid out systematically along transect lines formed by <1 m wide trails, and 14 were not located on trails but were chosen systematically along new routes. Given the narrow width and infrequent human use of trails, it is considered that this would not have overly biased results (Cornils *et al.* 2015). The distance between each point count was a minimum of 200 m, to minimise fly-between and double counting (Lloyd *et al.* 2000; Lee and Marsden 2008). Point counts, when possible, were conducted within the first 5 hours after dawn (Reid *et al.* 2012) and 3 hours before dusk (Cornils *et al.* 2015), when birds are most active (Poulsen and Lambert 2000), and were not conducted in rain, fog or high winds, as this reduces bird detectability and activity (Steadman and Freifeld 1998; Paz *et al.* 2013; Española *et al.* 2016).

GPS co-ordinates of all Negros Bleeding-heart sightings from both expeditions were mapped using ArcMap v.10.4 (ESRI 2016) along with the co-ordinates of habitat measurement points. The Negros Bleeding-heart was marked as associated to any habitat data-points within 60 m of a sighting’s co-ordinates, following protocol for the ground-forager dove *Leptotila wellsi* (Rivera-Milán *et al.* 2015). Four camera traps (Bushnell 12 MP Trophy Low Glow Essential HD Trail Camera) were placed at knee height above the ground from 21 June 2018 – 14 July 2018, and four more were added from 17 July until 6 June 2018. Camera traps were moved every 7-9 days (Rowcliffe *et al.* 2011) for a total of 161 camera-days. Camera traps were placed using purposive sampling to evenly cover the accessible areas and also focus on places where guides had seen the dove previously (Treves *et al.* 2010).

#### Measuring habitat

Ninety-three quadrats of 5 m radius were surveyed to measure habitat variables, a figure assumed to be representative of surrounding rainforest habitat (Peh *et al.* 2006; Posa and Sodhi 2006; Reid *et al.* 2012; Pangau-Adam *et al.* 2015). 64 of these quadrats were exactly coincident with point count locations and 13 with camera trap locations (figure 1). The remaining 16 did not coincide with another measurement point. All 93 quadrats were associated to any Negros Bleeding-heart sightings recorded within 60 m of them (Rivera-Milán *et al.* 2015), as stated in the bird survey methods. At each quadrat, GPS co-ordinates and an altitude reading were taken using a Garmin GPS unit (Garmin eTrex 30x), because altitude can be a major predictor of occurrence in forest bird species (Bibby 2000; Dallimer and King 2007; Paz *et al.* 2013). Tree Diameter at Breast Height (DBH) was measured in three categories: 0 to ≤25 cm (band 1), >25 to ≤50 cm (band 2), and >50 cm (band 3), following other bird-habitat studies (Paz *et al.* 2013; Zarones *et al.* 2013). Tree height was measured (Field Studies Council 2018) and branching architecture was noted following the guidelines in Bibby *et al.* (2000) to indicate recent forest disturbance levels, following which proportion of trees with a closed and regenerating forest structure was calculated. Canopy cover was estimated using a spherical densiometer (Forest Densiometers, Rapid City, ND) (Peh *et al.* 2006; Posa and Sodhi 2006; Reid *et al.* 2012), by calculating the mean canopy percentage cover from the north, east, south and west value. Percentage vegetation cover at ground level (<1 m) (“percentage ground cover”) and understorey cover at 1.5 m above the ground were estimated by eye by the first author (Posa and Sodhi 2006; Pangau-Adam *et al.* 2015). The presence or absence of any species of pandan *Pandanus sp.* and rattan (genus: *Calamus or Daemonorops*, Tesoro 2002) was noted (CCI report, personal communication). Distance to Sibaliw station was calculated using ArcMap, because it has been shown that species diversity can be higher closer to field stations (Campbell 2011).

### Data Analysis

#### Habitat preference

Data were analysed with a Generalised Linear Model (Zuur *et al.* 2013) in R v.3.6.1 (R Core Team 2019), in which the dependent variable was presence/absence of the Negros Bleeding-heart. Following Zuur *et al.* (2013), no outliers were found in the dataset. Multicollinearity was tested for using variance inflation factor (Zuur *et al.* 2013). Tree height was found to be highly correlated with DBH (R=0.67) using a pair plot (Zuur *et al.* 2013) hence was dropped from the model (O’Brien 2007; Zuur *et al.* 2010). All the explanatory variables were scaled to enable model convergence and effect size comparisons (Zuur *et al.* 2013; Abrahams *et al.* 2017). R package “MuMIn” version 1.43.6 (Bartoń 2015) was used for the GLM. A full model was created using the R function *glm*, following which automated model selection was done using the R function *dredge*, after which the most suitable selections were averaged using the R function *model.avg*, as described in Feld *et al.* (2016) and used in other bird-habitat studies (e.g. Xu *et al.* 2017; Lewis *et al.* 2018). The GLM was run using the following covariates: ground cover, understorey cover, canopy cover, altitude, proportion of trees with a closed forest branching structure, proportion of trees with a regenerating forest branching structure, presence of pandan, presence of rattan, distance to Sibaliw research station, and the number of trees in each of DBH band 1, 2 and 3 at each site.

#### Species distribution modelling

Negros Bleeding-heart distribution was modelled in MaxEnt v.3.4.1 (Phillips *et al.* 2017) for maximum entropy species distribution modelling (Merow *et al.* 2013; Morales *et al.* 2017). Two environmental predictor inputs were used: a map of tree cover at 30 m resolution (Hansen *et al.* 2013) and an elevation map at 90 m resolution (PhilGIS 2007), as the Negros Bleeding-heart is thought to be found in lowland forests up to 1000 m (BirdLife International 2017). While all 31 Negros Bleeding-heart sightings co-ordinates were inputted, due to the tree cover and elevation map resolutions, a sample size of 22 presence records was used for training. Feature types, selected automatically by MaxEnt, were hinge, linear and quadratic. The regularization parameter on MaxEnt was set to 2, which is a suitable figure to reduce over-fitting compared to the default setting of 1 (Radosavljevic and Anderson 2014); lower settings risk loss of predictive ability with independent test data, and higher settings can falsely mark areas as suitable (Radosavljevic and Anderson 2014). The background was cropped to include only the islands of the species’ known range (Merow *et al.* 2013), Panay and Negros. The model was run with 10,000 background points, which has been evaluated as high-performing number (Phillips and Dudik 2008) and used in studies on cryptic insect species (Zhao *et al.* 2019). Finally, the locations of Important Bird Areas (IBAs) (BirdLife International 2008b) were mapped over the species distribution modelling results to enable evaluation of conservation action.

## Results

The Negros Bleeding-heart was recorded 31 times in total. Camera traps recorded the Negros Bleeding-heart on three separate occasions (21/6/2018, 22/6/2018 and 24/6/2018) at the same location. Hunters were also recorded on two occasions along the same location.

### Habitat preference

All Negros Bleeding-heart sightings were associated with habitat plots measured (i.e. 31 out of the 93 quadrats). The full averaged GLM results (table 1) showed that understorey cover at 1.5 m above the ground was significant at the P <0.01 level (P = 0.001), showing a positive association with dense understorey cover. Canopy cover was significant at the P <0.05 level (P = 0.003), showing a positive association with dense canopy cover.

**Table 1.** Estimates of fixed effects of habitat variables on Negros Bleeding-heart presence or absence from Generalised Linear Model.

|  | Coefficient | Standard Error | z-value | p-value |
| --- | --- | --- | --- | --- |
| (Intercept) | -0.876 | 0.266 | 3.242 | 0.00119** |
| Ground cover (%) | -0.050 | 0.167 | 0.298 | 0.76591 |
| **Vegetation cover at 1.5 m above ground (%)** | 0.904 | 0.297 | 2.999 | **0.00271\*\*** |
| **Canopy cover (%)** | 0.778 | 0.367 | 2.091 | **0.03655\*** |
| Proportion of trees grown in closed canopy forest | -0.007 | 0.063 | 0.111 | 0.912 |
| Proportion of trees grown in open canopy or regenerating forest | 0.012 | 0.079 | 0.144 | 0.88571 |
| Pandan present | -0.005 | 0.051 | 0.097 | 0.92256 |
| Rattan present | 0.130 | 0.244 | 0.528 | 0.59721 |
| Altitude (m) | -0.007 | 0.062 | 0.115 | 0.90823 |
| Number of trees with a DBH $0 \leq 25$ cm | 0.054 | 0.170 | 0.317 | 0.75156 |
| Number of trees with a DBH $>25 \leq 50$ cm | -0.014 | 0.092 | 0.152 | 0.87929 |
| Number of trees with a DBH $>50$ cm | 0.226 | 0.274 | 0.818 | 0.41323 |

\*\*P <0.01, \*P <0.05

Coefficients and 95% confidence intervals of the explanatory variables in the GLM were graphed (figure 2).

**Figure 2.**
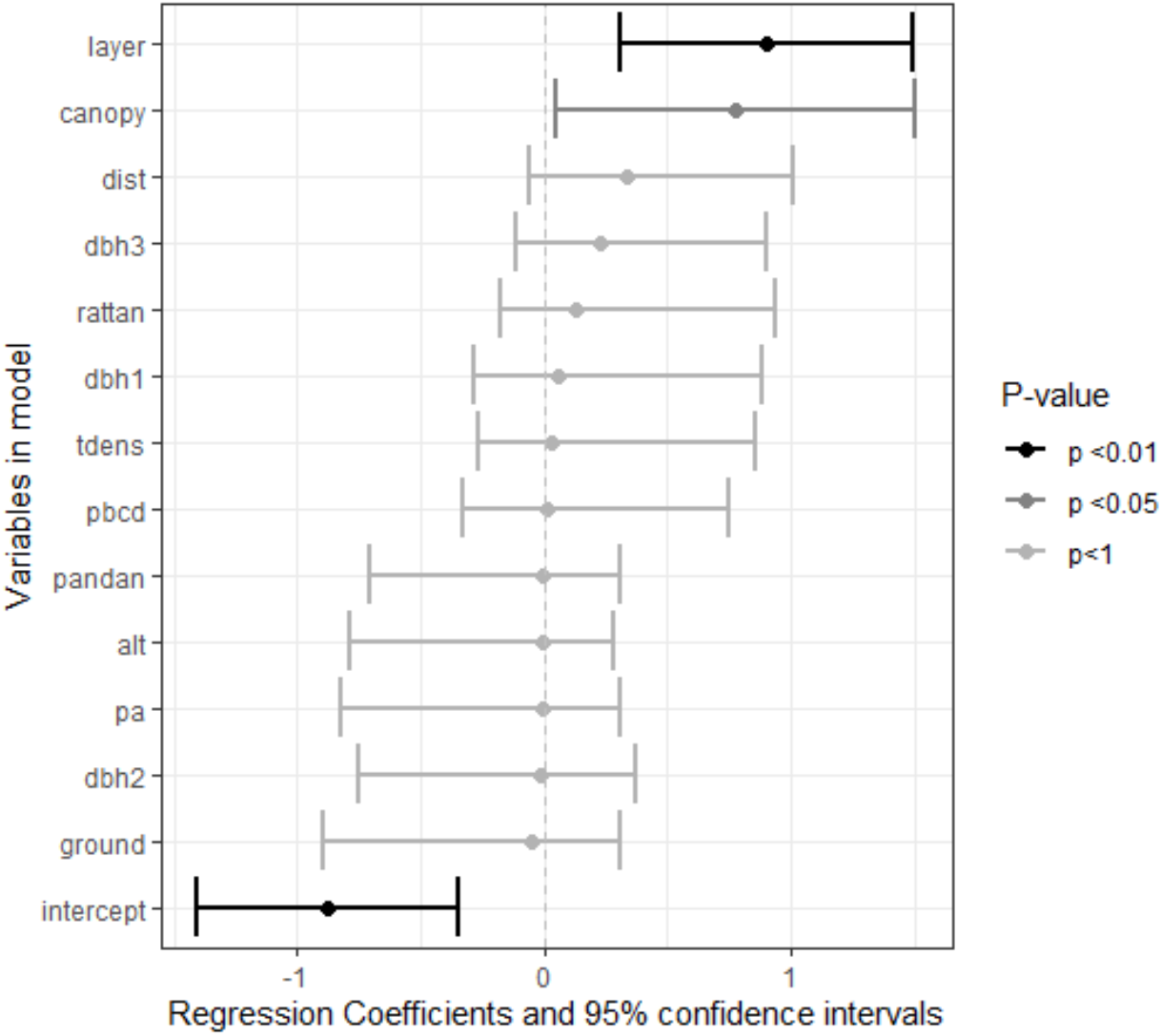
Coefficients and 95% confidence intervals of explanatory variables from Generalised Linear Model. Variable name abbreviations are as follows: layer = vegetation cover at 1.5 m above ground (%); canopy = canopy cover (%); dist = distance to Sibaliw research station (m); dbh1 = number of trees with a DBH 0 ≤ 25 cm; dbh2 =number of trees with a DBH >25 ≤ 50 cm; dbh3 = number of trees with a DBH >50 cm; ground = Ground cover (%); pa = proportion of trees grown in closed canopy forest; pbcd = proportion of trees grown in open canopy or regenerating forest; pandan = pandan presence, rattan = rattan presence, alt = altitude (m).

### Species distribution modelling

The species distribution modelling in MaxEnt produced probabilities of presence ranging from 0 to 1, with the most suitable habitat on Panay located around the Northwest Panay Peninsula Natural Park and the Central Panay Mountain Range, and the most suitable habitat in Negros found in the North Negros Natural Park, on the slopes of an area just outside the Mt Kanla-on Natural Park, and in an area to the south half-covered by Cuernos de Negros IBA (figure 3). However, very few areas are highly suitable, and those that exist appear fragmented, particularly on Negros. Furthermore, there are large unsuitable patches in high altitude areas due to this species’ preference for lower altitude forest at the centres of the North Negros Natural Park, Mount Kanla-on Natural Park, Cuernos de Negros and the Central Panay Mountain Range.

**Figure 3.**
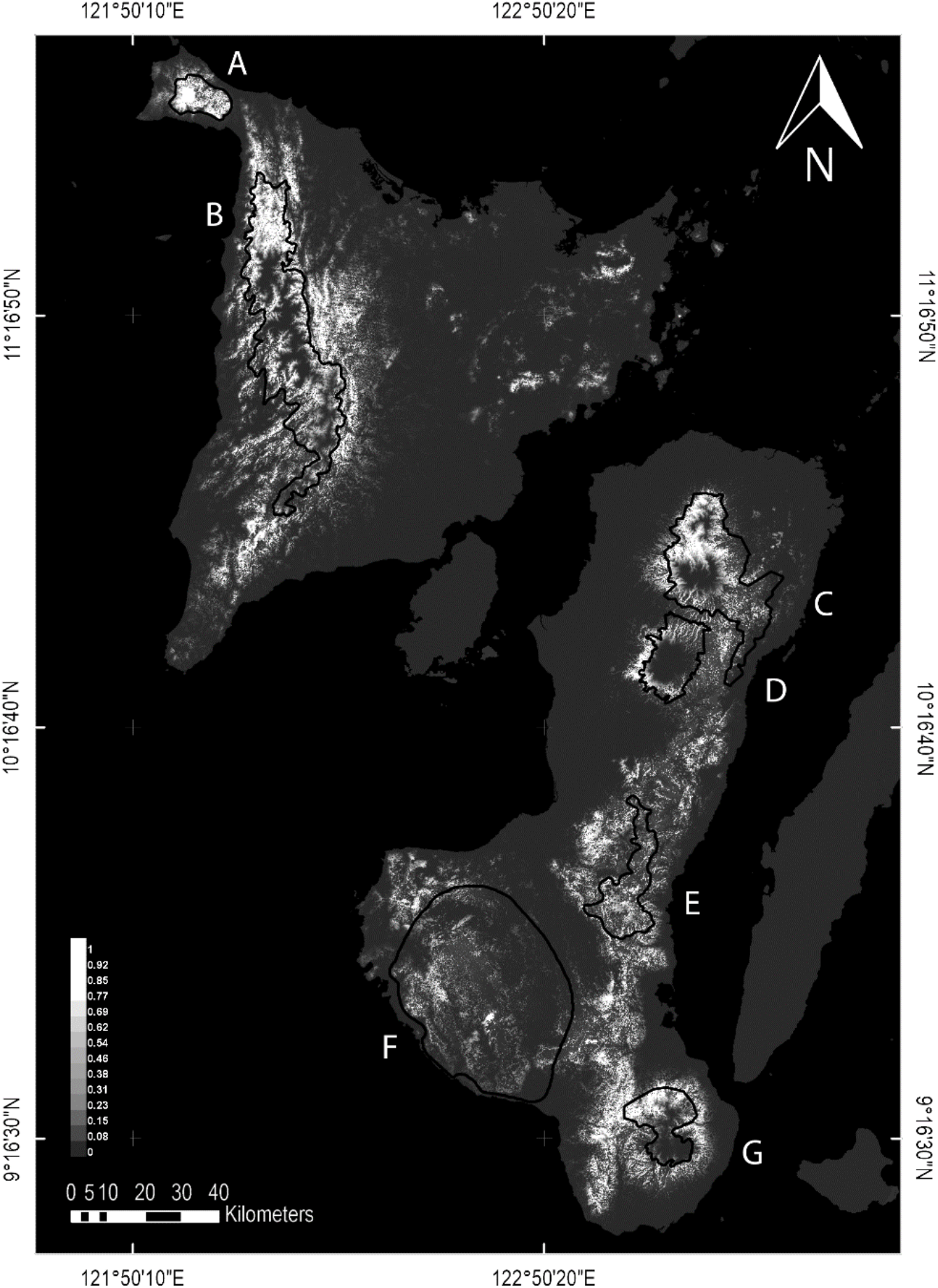
Species distribution modelling of the Negros Bleeding-heart on Panay and Negros. Scale is from white to dark grey, with probability of presence from 1 to 0 respectively. Locations of Important Bird Areas are shown as black outlines. On Panay: (A) Northwest Panay Peninsula Natural Park (BirdLife International 2018a), (B) Central Panay Mountain Range (BirdLife International 2018b). On Negros: (C) North Negros Natural Park (BirdLife International 2018c), (D) Mount Kanla-on Natural Park (BirdLife International 2018d), (E) Ban-ban (BirdLife International 2018e), (F) Southwestern Negros (BirdLife International 2018f), (G) Cuernos de Negros (BirdLife International 2018g). The white block on Panay shows the study site.

## Discussion

### Habitat preference

Understorey cover, mostly by herbaceous shrubs and ferns, was significantly associated with Negros Bleeding-heart presence. This association has been found for other bird species, postulated to be because it allows a balance between the ability to escape from predators and availability of attractive food resources (Lima and Dill 1990; Reid *et al.* 2004; Smith *et al.* 2017). The Luzon Bleeding-heart *Gallicolumba luzonica*, a close relative to the Negros Bleeding-heart, is reported to use thick undergrowth to escape predators (Del Hoyo *et al.* 1997), and the Negros Bleeding-heart itself is known to forage on the ground, tossing aside leaf litter in its search for food (Curio 2001). However, undergrowth cover could also relate to forest quality. Dense shrub or herb cover can be characteristic of secondary forests (Cochard *et al.* 2018), therefore suggesting that the Negros Bleeding-heart could be able to inhabit secondary forest, as found in other ground doves which favour primary forest (Blanvillain *et al.* 2002). However, the present study also idenfied a preference for closed canopy forest, which is characteristic of primary forest (Shoo *et al.* 2016). A preference for primary forest would be supported by numerous other studies showing that terrestrial birds like the Negros Bleeding-heart are often the species most affected by loss of primary forest (Sieving *et al.* 1996; Newmark *et al.* 2010; Powell *et al.* 2013; Bradfer-Lawrence *et al.* 2018) and are often unable to inhabit secondary forest (Peh *et al.* 2005; Stratford and Stouffer 2013). The ability of the Negros Bleeding-heart to use secondary forest remains unclear (BirdLife International 2017). While recorded in some secondary forest patches in this study, such areas within the Northwest Panay Peninsula Natural Park are small and surrounded by primary forest, and therefore the bird might move between patches to exploit resources but rely mainly on primary forest, as postulated by Peh *et al.* (2005) for bird species in Malaysian forest. Potential inability to use secondary forest could therefore explain why the dove is not found in some areas of secondary forest on Negros (BirdLife International 2017).

The habitat preference findings of this study are preliminary, and could be improved by a greater quadrat size for habitat measurement and more accurate linkage of point count results to habitat plots; while 60 m association distance has been used in other studies on similar species (Rivera-Milán *et al.* 2015), it adds uncertainty to the results. However, as little is known about this highly threatened species, the findings still have potential importance in directing conservation action. Research into Negros Bleeding-heart tolerance of secondary forest is strongly recommended, particularly considering how little primary forest remains across its range (BirdLife International 2017).

### Species distribution modelling

Species distribution modelling shows that the areas of suitable habitat are those which are forested and at low altitude, as previously reported to be requirements of the Negros Bleeding-heart (BirdLife International 2017). The model also highlights that large proportions of the core areas inside many IBAs and protected areas maybe less suitable than previously thought, particularly within the protected areas of the North Negros Natural Park and Mt Kanla-on Natural Park, and the unprotected IBAs of the Central Panay Mountains and Cuernos de Negros, with some edge areas appearing more suitable than within the core area. This is a particular concern because the edges of reserves are often subject to abiotic edge effects that can reduce habitat quality, particularly for understorey birds (Pohlman *et al.* 2007; Neate-Clegg *et al.* 2016), and to greater human pressures, such as encroachment for agriculture, charcoal logging or hunting by those living on the borders (Pedregosa-Hospodarsky *et al.* 2009).

The North Negros Natural Park contains one of the only remaining lowland forest sites on Negros, although it is largely secondary forest (BirdLife International 2018c), and species distribution modelling shows a significant amount of potentially suitable habitat near the eastern border of the park. The park reportedly contains the Negros Bleeding-heart (BirdLife International 2018c). However, the species is not mentioned in an Endangered bird biodiversity survey undertaken there in 2002 (Hamann 2002), and nor was it observed during a 20 day study there in May 2009 (Pedregosa-Hospodarsky *et al.* 2009), the same month in which the present study recorded 28 sightings in the Northwest Panay Peninsula Natural Park. In addition, despite active protection, North Negros Natural Park is under pressure in almost every municipality within it from illegal logging and hunting (including using snares that catch ground-dwelling birds like the Negros Bleeding-heart) (Pedregosa-Hospodarsky *et al.* 2009). Therefore, rapid conservation action is required.

Species distribution modelling shows that there is a small area of suitable habitat for the Negros Bleeding-heart along the boundary of Mt Kanla-on Natural Park. One Negros Bleeding-heart was seen on Mt Kanla-on at 900 m in 1992 (Brooks *et al.* 1992), but a 2007 survey there undertaking field surveys and interviews failed to encounter it (Cariño, 2007). Furthermore, the Negros Bleeding-heart is thought to have an an altitudinal limit of 1,000 m (BirdLife International 2017), a lower altitude than most forest around Mt Kanla-on (Brooks *et al.* 1992), hence it seems probable that the bird is no longer found within the park. Any populations just outside the boundary would be threatened by the continued deforestation on the lower slopes (BirdLife International 2018d), demonstrating that Mt Kanla-on Natural Park may not be the best placed to protect the dove.

Of the non-protected IBA sites on Negros, the largest indicated area of potentially suitable habitat for the Negros Bleeding-heart is around the Cuernos de Negros IBA, half within the reserve and the rest just outside the borders. Cariño (2007) recorded the species several times in Siaton, neighbouring the border, and it was reported inside the IBA eight times aurally and once visually in the CCI report (personal communication) during their 2016 survey. However, the CCI survey duration was 4 days, and another 9-week survey in the same area in 2017, using camera traps, line transects and point counts, failed to encounter the Negros Bleeding-heart (Cantero-Sanchez 2018). Furthermore, Bristol Zoological Society have studied the wider Cuernos de Negros IBA for 4 years rom 2014-2017, obtaining a minimum of 347 camera trap nights, and the Negros Bleeding-heart was never sighted (Falcidia 2017). It is therefore possible that the dove may not be in high density or even present in the Cuernos de Negros IBA, given the high altitude of this area. Hence, this may not be a priority site on which to focus Negros Bleeding-heart protection action.

Of the two IBAs in Panay, it is clear that the Negros Bleeding-heart is present in the Northwest Panay Peninsula Natural Park. From the number of sightings obtained in this study compared to other recent studies on Negros, and from the species distribution modelling results, it also appears to be the most effective existing protected area on Panay and Negros for the Negros Bleeding-heart. It is therefore a critical stronghold for the species, requiring continued conservation action, which is ongoing (PhilinCon 2018). Furthermore, there appear to be large areas of potentially suitable habitat for the Negros Bleeding-heart around and within the Central Panay Mountain Range, although this is not yet a formal protected area (Haribon Foundation 2016). While the Negros Bleeding-heart is reported to exist in the Central Panay Mountain Range by several sources (De Soye 1997; Pedregosa-Hospodarsky 2008; Manila Times 2010), to our knowledge, there is no recent scientific survey report. Therefore, this area is a priority for survey effort, particularly considering the likely extreme rarity of the Negros Bleeding-heart on Negros.

Finally, species distribution modelling shows that the suitable habitat patches throughout Panay and Negros are not well linked, which is detrimental to maintaining small populations (Kramer *et al.* 2009; Spigler *et al.* 2017; Gómez-Sánchez *et al.* 2018). *Ex situ* conservation is a useful support in these situations (Braverman 2014). Captive breeding has begun for the Negros Bleeding-heart at the Centre for Tropical Conservation Studies in Negros, where the captive population now numbers 18 birds (EDGE of Existence 2019) and it is recommended that this work continues alongside *in situ* conservation measures.

## Conclusion

The Negros Bleeding-heart has a preference for dense understorey vegetation and dense canopy cover, and it currently seems that the largest area of suitable habitat is in the Northwest Panay Peninsula Natural Park on Panay. On Negros, Mt Kanla-on Natural Park may not contain the best habitat for the Negros Bleeding-heart due to its altitude, and further protection of the North Negros Natural Park eastern border is needed. It is essential that laws against deforestation and hunting in all remaining population refuges are strictly enforced in order to halt the decline in numbers. In addition, it is recommended that future research effort focuses on two areas: to establish the degree to which the Negros Bleeding-heart requires primary forest rather than secondary, in order to direct survey and conservation action, and to assess its population status in the Central Panay Mountain Range and the North Negros Natural Park, which have the potential to be suitable sites for Negros Bleeding-heart conservation.

## Acknowledgements

The authors thank everyone at PhilinCon for making this study possible: guides Potpot Fernandez and Jun Tacud, Rhea Santillan for helping organise the logistics, the Sibaliw research station staff and porters, including Momel, Ulysses and Dudong, and Professor Dr. Eberhard Curio, PhilinCon President.

